# An open source statistical and data processing toolbox for wide-field optical imaging in mice

**DOI:** 10.1101/2021.04.07.438885

**Authors:** Lindsey M. Brier, Joseph P. Culver

## Abstract

Wide-field optical imaging (WOI) produces concurrent hemodynamic and cell-specific calcium recordings across the entire cerebral cortex. There have been multiple studies using WOI to image mouse models with various environmental or genetic manipulations to understand various diseases. Despite the obvious utility of pursuing mouse WOI alongside human functional magnetic resonance imaging (fMRI), and the multitude of analysis toolboxes in the fMRI literature, there is not an available open-source, user-friendly data processing and statistical analysis toolbox for WOI data. Here, we present our MATLAB toolbox for pre-processing WOI data, as described and adapted to combine processing techniques from multiple WOI groups. Additionally, we provide multiple data analysis packages and translate two commonly used statistical approaches from the fMRI literature to the WOI data. To illustrate the utility, we demonstrate the ability of the processing and analysis framework to detect a well-established deficit in a mouse model of stroke. Additionally, we evaluate resting state data in healthy mice.

## Introduction

Functional neuroimaging has enhanced our study of systems neuroscience and understanding of neural networks^1,2^. Mainly, this has been accomplished with blood oxygen level dependent (BOLD) fluctuations in functional magnetic resonance imaging (fMRI) data in human subjects^3^. In order to better understand human conditions, there has been an increase in functional neuroimaging in animal models, also performed using fMRI^4–6^. However, the relatively small size of the mouse brain offers multiple technical and logistical challenges with fMRI. Therefore, there has been a transition to using wide-field optical imaging (WOI) techniques in the mouse, yielding similar blood-based surrogates of neural activity at a similar spatial scale^7^. The advent of genetically encoded calcium indicators (GECIs) enables cell-specific labelling and increased temporal resolution compared to traditionally measured hemodynamics^8–10^. Combined hemoglobin and fluorophore imaging is readily available with optical imaging systems and harnesses the advantages of GECIs as well as maintains a translatable blood-based recording directly comparable to human fMRI. WOI analysis faces many of the same procedural steps and therefore difficulties as those experienced with fMRI analysis, such as data processing, visualization, and statistical testing. However, the relative novelty of WOI compared to fMRI means that there is a need for many of the solutions within the fMRI community to be translated to the WOI data analysis communities.

One of the biggest statistical challenges within the functional neuroimaging community is the problem of correcting for multiple statistical tests. Many solutions have been proposed within the fMRI community^11,12^, however, in general these have not been translated into an easy-to-use WOI toolbox. Historically, functional connectivity (FC) is examined using both a seed-based and a matrix-based approach^13^. For seed-based maps, common practice includes performing a pixel or voxel-wise statistical test (e.g., student’s t) resulting in thousands of tests being performed within the field-of-view (FOV). The most stringent correction (i.e., Bonferroni), assumes each statistical test is independent^14^. This is certainly not the case when examining neighboring pixels within a brain region for multiple reasons. For most mesoscopic WOI instruments, the full-width half-maximum (FWHM) spans multiple pixels thus rendering each pixel not independent from an instrumentation point of view. Additionally, from a biological point of view, it is reasonable to assume an amount of dependence between neighboring pixels within the same brain region. A more plausible approach borrowed from fMRI to handle the multiple comparisons problem uses a clustering analysis, coupled with random field theory, to weight larger regions of interest (ROI, i.e., large clusters) of contiguous neighboring significant pixels as more likely to be a statistically significant finding than small ROI’s^15,16^. However, this approach has not been translated from fMRI voxel space to WOI pixel space application.

Although FC matrices inherently go through a certain amount of data reduction, a well-sampled FOV can still lead to hundreds or thousands of multiple comparisons. A simple data reduction technique that combines the average FC value within an entire cortical area (e.g., as determined by the Paxinos atlas^7^) into a singular network value can be more informative and provide stronger statistical power. Network level analysis, the fMRI equivalent of enrichment analysis in the genome-wide association studies (GWAS) literature, seeks to highlight networks significantly correlated with a singular value (e.g., other networks, behavior, clinical outcome), rather than each individual cortical connection^17^. Once the data has been reduced to networks, then a Bonferroni correction can be applied on the remaining elements.

Here, we provide a mouse optical data processing toolbox to streamline and make pre-processing steps transparent and user friendly. Within it, we adopt the fMRI cluster size-based approach to determining statistical significance to wide-field optical FC mapping in the context of photothrombotic stroke. Additionally, we provide our methods in data reduction and matrix enrichment testing to be used throughout the WOI community.

## Methods

### Animals

For bilateral FC analysis, four 3-4 month old mice (2 male, 2 female) were imaged at baseline (Day 0) and on Day 3 post photothrombotic stroke to left somatosensory forepaw cortex. For FC matrix analysis, a total of 16 female (ranging from 3-7 months old) mice were used. All mice (N=20) were *Thy1*-GCaMP6f (Jackson Laboratories Strain: C57BL/6J-*Tg(Thy1-GCaMP6f)GP5*.*5Dkim*; stock: 024276). These mice express the protein GCaMP6f in excitatory neurons, primarily in cortical layers ii, iii, v, and vi^8^. All studies were approved by the Washington University School of Medicine Animals Studies Committee and follow the guidelines of the National Institutes of Health’s Guide for the Care and Use of Laboratory Animals.

### Surgical preparations

Prior to imaging, typical surgical preparations were implemented^9,18^. Briefly, an optically transparent plexiglass window was implanted with translucent dental cement (C&B-Metabond, Parkell Inc., Edgewood, New York) following a midline incision and clearing of skin and periosteal membranes. The window covered the majority of the dorsal cortical surface and provided an anchor for head fixation and allowed for chronic, repeatable imaging. The N=16 mice used for FC matrix computation had stainless steel EEG self-tapping screws (BASI Inc., West Lafayette, IN, USA) fixed at approximately -1mm posterior to bregma, and +/- 5mm lateral to bregma (near barrel/auditory cortex), although EEG recording was omitted for the present report. The dataset used in the following analyses consists of two ten-minute imaging runs from each mouse in the N=16 group. One ten-minute run from one mouse was discarded due to increased variance in the raw light levels.

### Photothrombosis

In N=4 (2 male, 2 female, all are mice that did not receive EEG screws), mice were secured in a stereotaxic frame under isoflurane anesthesia. 200 *μl* of Rose Bengal (Sigma Aldrich) dissolved in saline (10 g/liter) was injected intraperitoneally. After 4 minutes, a 532-nm diode-pumped solid-state laser (Shanghai Laser & Optics Century) was focused to 2.2mm left and 0.5mm anterior to bregma with a 0.5mm spot size and at 23mW for 10 minutes^19^. Mice were imaged at baseline (i.e., prior to photothrombosis (Day 0)), and 72 hours post (Day 3). The dataset used in the following analyses consists of two five-minute imaging runs from each mouse.

### Fluorescence and Optical Intrinsic Signal (OIS) imaging

Mice were head-fixed in a stereotaxic frame and body secured in a black felt pouch for imaging. Sequentially firing LEDs (Mightex Systems, Pleasanton California) passed through a series of dichroic lenses (Semrock, Rochester New York) into a liquid light guide (Mightex Systems, Pleasanton California) that terminated in a 75mm f/1.8 lens (Navitar, Rochester New York) to focus the light onto the dorsal cortical surface. LEDs consisted of 470nm (GCaMP6f excitation), 530nm, 590nm, and 625nm light. An sCMOS camera (Zyla 5.5, Andor Technologies, Belfast, Northern Ireland, United Kingdom) coupled to an 85mm f/1.4 camera lens (Rokinon, New York New York) was used to capture fluorescence/reflectance produced at 16.8 Hz per wavelength of LED. A 515nm longpass filter (Semrock, Rochester New York) was used to discard GCaMP6f excitation light. Cross polarization (Adorama, New York New York) between the illumination lens and collection lens discarded artifacts due to specular reflection. The field-of-view (FOV) recorded covered the majority of the convexity of the cerebral cortex (∼1.1cm^2^), extending from the olfactory bulb to the superior colliculus. All imaging data were binned in 156×156 pixel^2^ images at approximately 100 *µ*m^2^ per pixel.

### Imaging data processing

Image processing followed methods previously described^8,20^ and summarized here. Images were spatially downsampled to 78×78 pixel^2^ and a frame of ambient baseline light levels was subtracted from the time series data. Data were temporally downsampled by a factor of 2 and then spatially and temporally detrended. The logarithmic mean of frames corresponding to reflection data produced by the 530nm, 590nm, and 625nm light were used to solve the modified Beer Lambert law to yield fluctuations in oxygenated and deoxygenated hemoglobin. Frames corresponding to fluorescence data were mean normalized and corrected by approximating hemoglobin absorption of the excitation and emission light, following which the logarithmic mean was taken. All data were spatially smoothed with a 5×5 Gaussian filter. The global signal was regressed from the time series data, data were Affine transformed to common Paxinos atlas space and data were filtered with a 0.4-4.0Hz Butterworth bandpass filter.

### Functional connectivity

Functional connectivity (FC) analysis refers to the calculation of a Pearson correlation coefficient between two time traces:

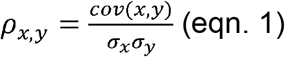

where ρ_*x,y*_ is the Pearson correlation coefficient between time traces *x* and *y*, *cov(x,y)* is the covariance between time traces *x* and *y*, and σ_*x*_, σ_*y*_are the standard deviations within time trace *x* and time trace *y*, respectively. For bilateral FC maps, the Pearson correlation coefficient is computed between pixels on the left-hand side of the brain and the respective mirror imaged pixel on the right-hand side. For FC matrices, the Pearson correlation between the average time trace within two seed regions is computed.

### Cluster-based thresholding

A challenge with analyzing the statistical significance in functional imaging is managing the multiple comparisons problem. Here, we used a cluster size-based method that leverages the spatial connection between pixels, and credits large clusters as having more statistical significance than small clusters with the same peak t-value. More specifically, we used a cluster-size based thresholding method (determined by the cluster-size limit, *k*_*α*_)to analyze two-dimensional FC maps, and to ensure the family-wise error (FWE) rate did not exceed 5%. A random field theory (RFT) approach was adapted from the fMRI and DOT literature^15,16^. Using RFT, we are able to approximate the expected number of clusters (*m*) in an image at a given z-score threshold (*Z*_*t*_, Figure 2A):

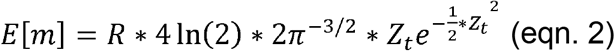

Which holds true for increasing values of *Z*_*t*_. Here, we use *Z*_*t*_ =3.09 which corresponds to a false positive rate of 0.001 at the pixel level. *R* represents the number of resolution elements provided by the optical system:

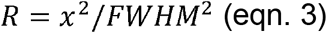

Where *x* is the number of pixels in the image in one dimension and *FWHM* is the full width half maximum of the point spread function estimated from the spatial autocorrelation of a fully processed image in one dimension. Here, we use *x*=78 pixels and calculate a *FWHM* of about 14 pixels. We are able to then approximate the expected number of pixels (*N*) above *Z*_*t*_ by:

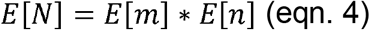

Where *n* is the pixel count within a cluster and:

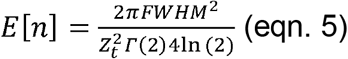

where Г is the gamma function. The probability that *n* will exceed any threshold, *x*, can be modeled by the exponential function:

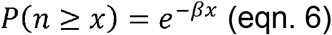

Where *β* can be expressed as:

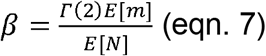

Substituting eqn. 5 into eqn. 4 and then the newly formed eqn. 4 into eqn. 7 rearranges the above to:

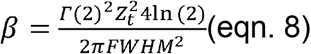

Providing an equation (with all previously solved for values) that varies with the inverse of the squared *FWHM*. We only want clusters to survive thresholding with a family wise error rate of 0.05 (*α*), therefore, we want to find the pixel count threshold, *k*_*α*,_ that would result in at least one cluster surviving threshold when there is no true significant difference, 5% of the time. Essentially, we are asking for a *k*_*α*_ where the following is true:

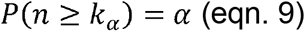

Assuming no clusters are a true positive. Which, mathematically speaking, is the same as solving for 1 minus the probability that no clusters have a pixel count above threshold *k*_*α*_ (Figure 2B):

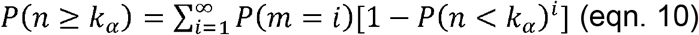

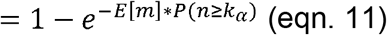

Which yields:

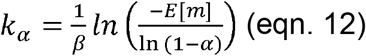

After substituting eqn. 6 into eqn. 11 and solving for *k*_*α*_.

Pixel-wise t-tests were performed to compare Day 0 and Day 3 stroke bilateral FC maps. These t-test maps were then thresholded, leaving pixels with a t-value corresponding to p<0.05. Any clusters of surviving pixels with a pixel count greater than *k*_*α*_ survived significance testing according to this cluster size-based technique.

### Matrix enrichment

The challenge of managing the multiple comparisons problem is still present with FC matrix analysis. Additionally, the non-intuitive presentation of such analysis precludes interpretation to non-imaging groups. Therefore, we have established a simple solution to both problems. The average intra-cortical region Pearson correlation coefficient is stored in the same surface area as presented in the individual seed method^17^. This data reduction method can increase the maximum Bonferroni p-value threshold by an order of magnitude as well as increase the readability of the plots.

### Data and code availability

In order to promote validation and comparative analyses by external groups, data and specific code will be made available through requests to the corresponding author. The toolbox presented here (for WOI pre-processing and analysis), instructions, and example excel file is available online (https://github.com/brierl/Mouse_WOI) and was used for all present analysis. Each major wrapper and respective subroutines are detailed in Tables 1-8. Example data is available on Figshare (https://figshare.com/articles/dataset/Mouse_WOI_Test_Data/14204549) and detailed in Table 9.

**Table 1:** Functions for all wrappers (wrappers as named on the GitHub repository)

| Function | Description |
| --- | --- |
| excel_reader.m | Read in mouse specific information |
| image_system_info.m | Read in optical system specific information |

## Results

### Toolbox overview

We seek to distribute a comprehensive, easy-to-use, open-source toolbox for mouse WOI data processing and analysis. Within this toolbox, we have translated multiple techniques from the human fMRI literature to the WOI mouse world. The toolbox is largely split up into three pipelines (Figure 1, Tables 1-8). User input is needed to organize mouse and instrument specific details to be read in (Table 1), along with a single imaging frame (Figure 1A). That single frame is used for the creation of mask and seed files. Binary masks are created using the roipoly.m function in MATLAB and the user traces the perimeter of the brain visible in the FOV. Seed regions (for affine transform) are created by the user identifying the anterior suture and lambda. Based on user input in Figure 1A, pre-processing of the data is automatically performed following the steps outlined in Figure 1B. Instrumentation details from Figure 1A will allow the program to split up the imaging frames to solve for either hemoglobin, fluorophore, or both concentration changes over time. All these steps are organized into independent sub-routines, allowing the user to add or subtract steps within the master scripts (e.g., Proc1_sys_dep.m, Proc2.m, Table 2) as needed. There are optional quality control (QC) outputs before and after the data is optionally temporally filtered that can be used for data scrubbing procedures (under development). After pre-processing of the data, the analysis pipelines are largely similar (Figure 1C, Tables 3-8). First, the analysis is performed on each individual run for each individual mouse, followed by an averaging across mice. A statistical comparison is then performed on the data (user specified) and the cluster size-based thresholding is used to determine regions of statistical significance. Only with the seed-based FC analysis are matrices produced and does matrix enrichment occur. The available analysis pipelines include: Seed-wise FC (Table 3, Figure 3), bilateral FC (Table 4, Figure 2C), SVR multivariate FC^21^ (Table 5), spectral topography^9^ (Table 6), a neurovascular coupling (NVC) approximation^8^ (Table 7), and node degree FC^22^ (Table 8).

**Table 2:**
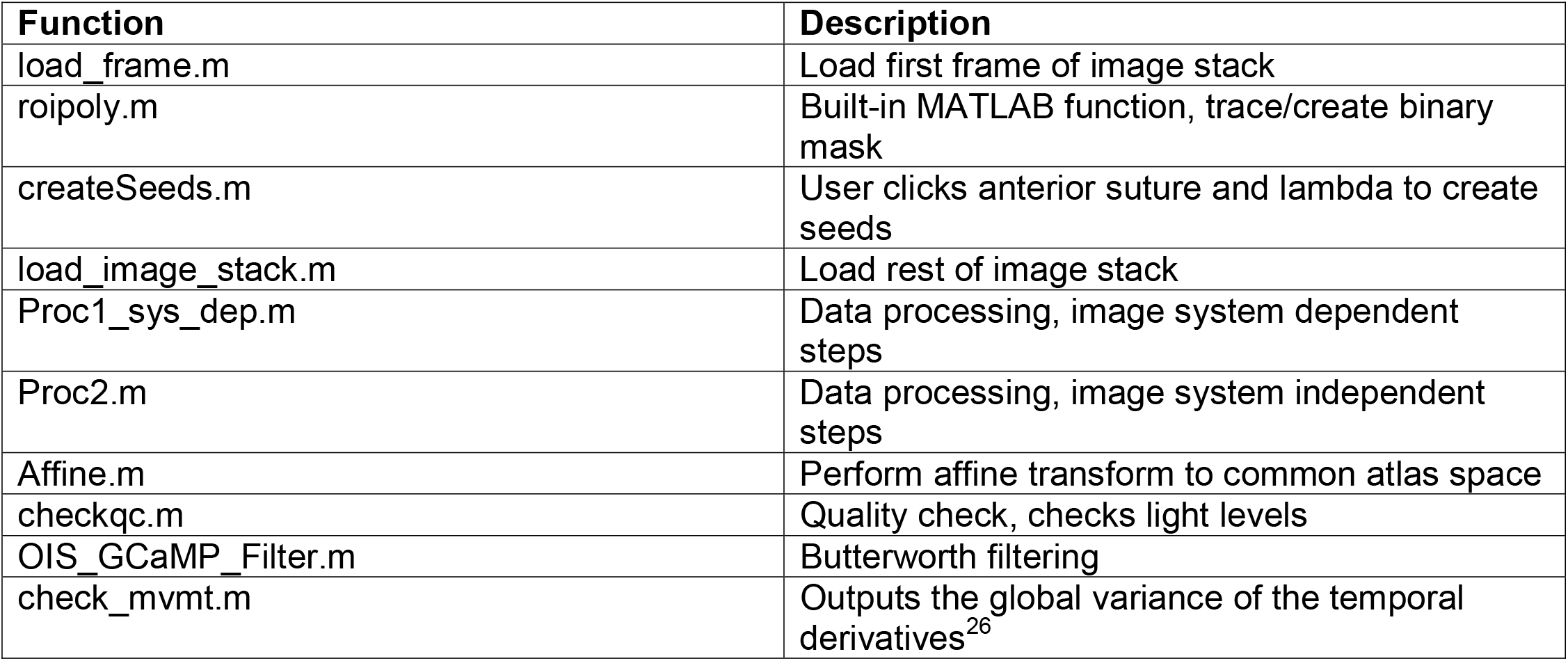
Data processing subroutines (Wrapper: MOUSE_MASTER_Proc)

| Function | Description |
| --- | --- |
| load_frame.m | Load first frame of image stack |
| roipoly.m | Built-in MATLAB function, trace/create binary mask |
| createSeeds.m | User clicks anterior suture and lambda to create seeds |
| load_image_stack.m | Load rest of image stack |
| Proc1_sys_dep.m | Data processing, image system dependent steps |
| Proc2.m | Data processing, image system independent steps |
| Affine.m | Perform affine transform to common atlas space |
| checkqc.m | Quality check, checks light levels |
| OIS_GCaMP_Filter.m | Butterworth filtering |
| check_mvmt.m | Outputs the global variance of the temporal derivatives<sup>26</sup> |

**Table 3:** Seed-wise FC subroutines (Wrapper: MOUSE_MASTER_FC)

| Function | Description |
| --- | --- |
| calc_fc.m | Calculates Pearson correlation per seed set per run |
| visualize_fc.m | Visualize seed-wise FC maps per run |
| visualize_fc_matrix.m | Visualize FC matrices per run |
| FC_AVG.m | Average FC runs across mice |
| visualize_fc_avg.m | Visualize averaged FC maps across mice |
| visualize_fc_avg_matrix.m | Visualize averaged FC matrices across mice |
| cluster_threshold.m | Solve for cluster-size based threshold |
| FC_ttest.m | Perform pixel-wise t-tests on FC maps |
| visualize_fc_ttest.m | Visualize results from t-tests on FC maps |
| FC_Matrix_ttest.m | Perform comparison-wise t-tests on FC matrices |
| visualize_fc_ttest_matrix.m | Visualize results from t-tests on FC matrices |
| prep_matrix_simple.m | Perform matrix enrichment |
| visualize_fc_e_matrix.m | Visualize enriched FC matrices |
| FC_e_Matrix_ttest.m | Perform comparison-wise t-tests on enriched FC matrices |
| visualize_fc_ttest_e_matrix.m | Visualize results from t-tests on enriched FC matrices |

**Table 4:**
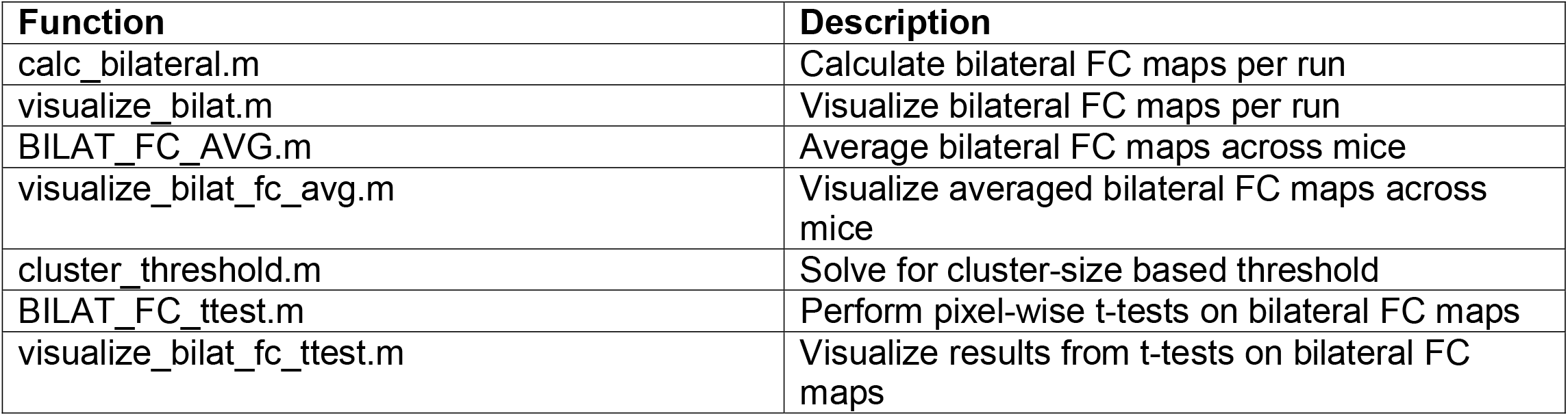
Bilateral FC subroutines (Wrapper: MOUSE_MASTER_Bilat)

| Function | Description |
| --- | --- |
| calc_bilateral.m | Calculate bilateral FC maps per run |
| visualize_bilat.m | Visualize bilateral FC maps per run |
| BILAT_FC_AVG.m | Average bilateral FC maps across mice |
| visualize_bilat_fc_avg.m | Visualize averaged bilateral FC maps across mice |
| cluster_threshold.m | Solve for cluster-size based threshold |
| BILAT_FC_ttest.m | Perform pixel-wise t-tests on bilateral FC maps |
| visualize_bilat_fc_ttest.m | Visualize results from t-tests on bilateral FC maps |

**Table 5:**
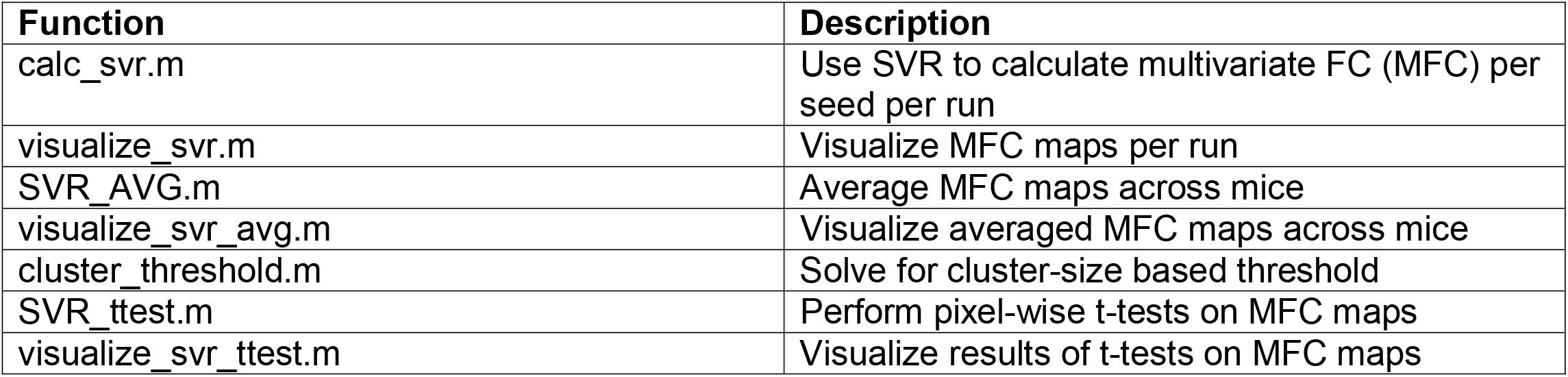
SVR multivariate FC subroutines (Wrapper: MOUSE_MASTER_SVR)

| Function | Description |
| --- | --- |
| calc_svr.m | Use SVR to calculate multivariate FC (MFC) per seed per run |
| visualize_svr.m | Visualize MFC maps per run |
| SVR_AVG.m | Average MFC maps across mice |
| visualize_svr_avg.m | Visualize averaged MFC maps across mice |
| cluster_threshold.m | Solve for cluster-size based threshold |
| SVR_ttest.m | Perform pixel-wise t-tests on MFC maps |
| visualize_svr_ttest.m | Visualize results of t-tests on MFC maps |

**Table 6:**
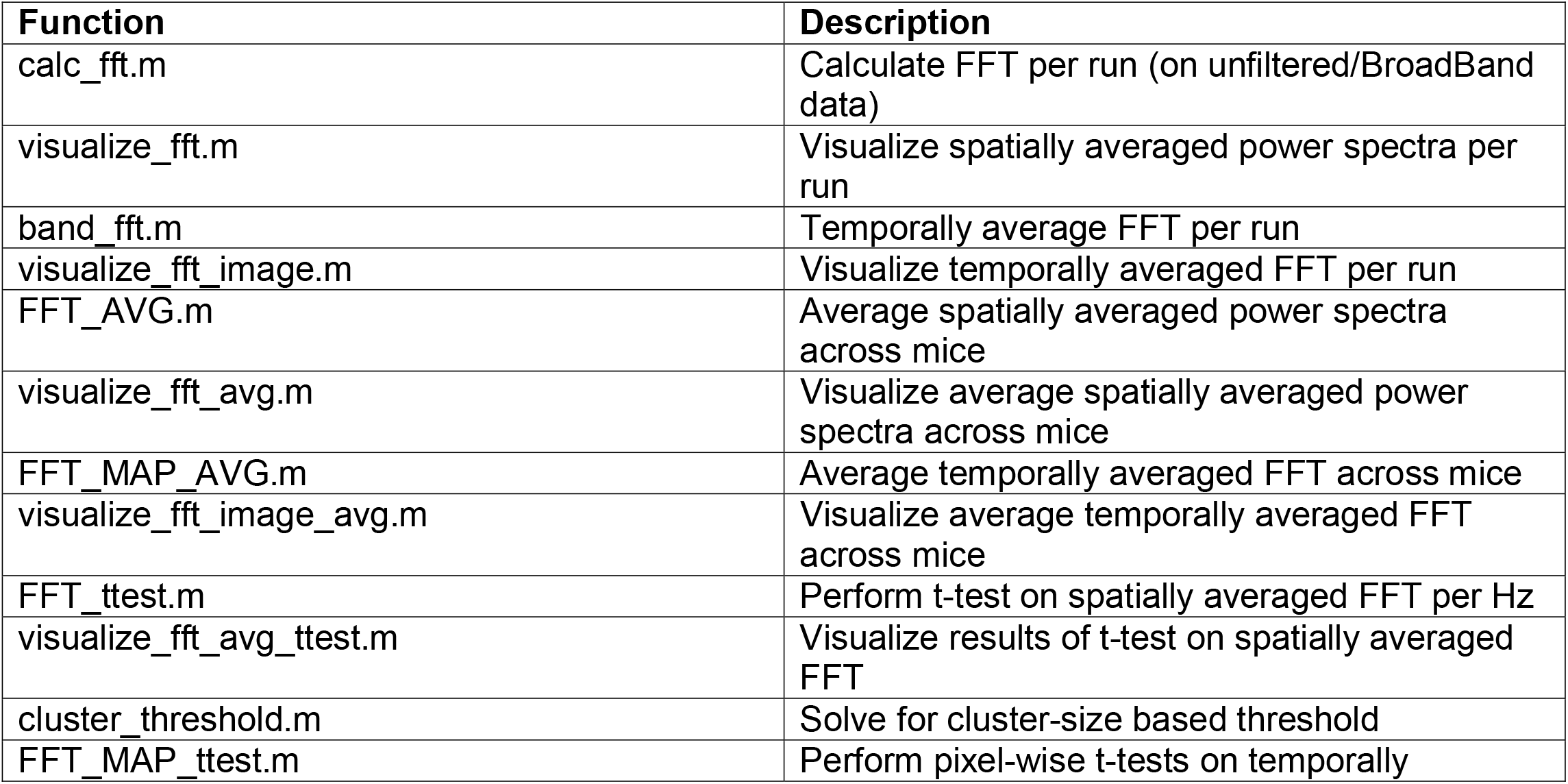
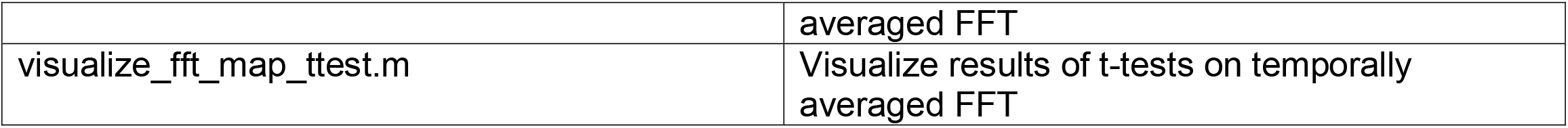
Spectral subroutines (Wrapper: MOUSE_MASTER_Spectral)

**Table 7:** NVC subroutines (Wrapper: MOUSE_MASTER_NVC)

| Function | Description |
| --- | --- |
| NVC_bypixel.m | Calculate neurovascular coupling approximation per run |
| visualize_nvc.m | Visualize neurovascular coupling approximation per run |
| AVG_NVC.m | Average NVC across mice |
| visualize_nvc_avg.m | Visualize averaged NVC across mice |
| cluster_threshold.m | Solve for cluster-size based threshold |
| NVC_ttest.m | Perform pixel-wise t-tests on NVC |
| visualize_nvc_ttest.m | Visualize results of t-tests on NVC |

**Table 8:** Node degree FC subroutines (Wrapper: MOUSE_MASTER_Node)

| Function | Description |
| --- | --- |
| calc_fc_node.m | Calculate node degree per run |
| visualize_nodes.m | Visualize node degree per run |
| FC_Node_AVG.m | Average node degree across mice |
| visualize_fc_node_avg.m | Visualize averaged node degree across mice |
| cluster_threshold.m | Solve for cluster-size based threshold |
| FC_Node_ttest.m | Perform pixel-wise t-tests on node degree |
| visualize_fc_node_ttest.m | Visualize results of t-tests on node degree |

**Table 9:** Sample data available on FigShare

| Type | Dataset |
| --- | --- |
| pixels x pixels x frames, resting state, 10 min | 190930-1185-LPS5Hr8Ms1-fc1.mat |
| pixels x pixels x frames, resting state, 10 min | 190930-1185-LPS5Hr8Ms1-fc2.mat |
| pixels x pixels x frames, resting state, 10 min | 190930-1186-LPS5Hr8Ms1-fc1.mat |

**Figure 1:**
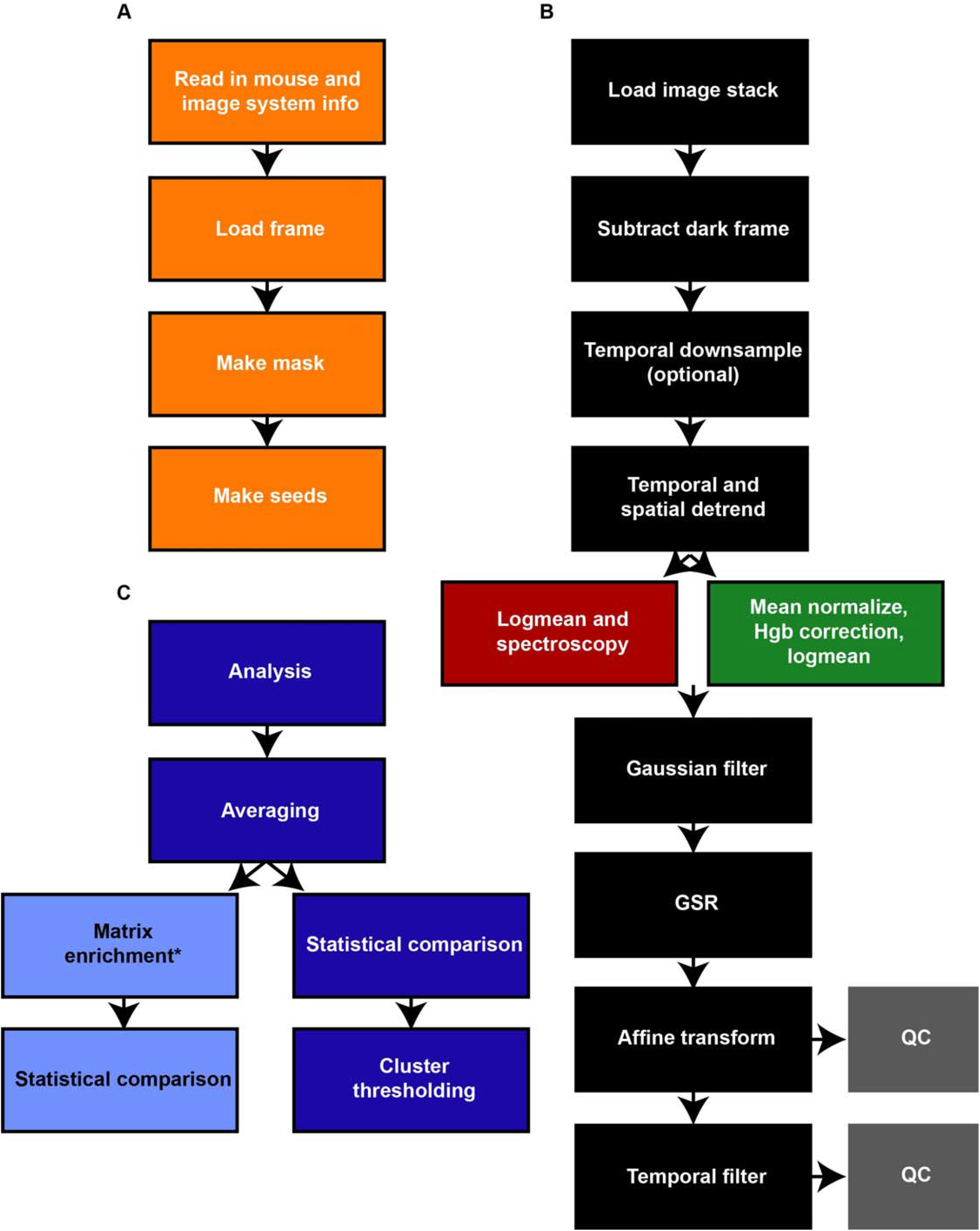
WOI mouse toolbox flowchart. A) User input needed for these steps. Excel sheet updated with mouse information as well as image_system_info.m updated with image system information is read into the pipeline. A single frame is loaded and a user defined binary mask (using the roipoly.m function) and seed locations are created. B) These steps will run without user input. Step-wise illustration of the data pre-processing pipeline color-coded for hemoglobin (red), fluorophore (green), or both (black, grey) specific steps. C) After the pre-processing steps in A,B), the toolbox supports various types of data analysis (e.g., seed-wise FC, bilateral FC, node degree FC, spectral) and all follow the dark blue illustrated analysis pipeline. The initial analysis is performed per run, per mouse, followed by averaging across mice. A pixel-wise statistical comparison is performed (e.g., t-test) and the cluster size-based thresholding is implemented to correct for multiple comparisons. *Only for seed-wise FC are matrices created and fed through the enrichment pipeline (light blue).

**Figure 2:**
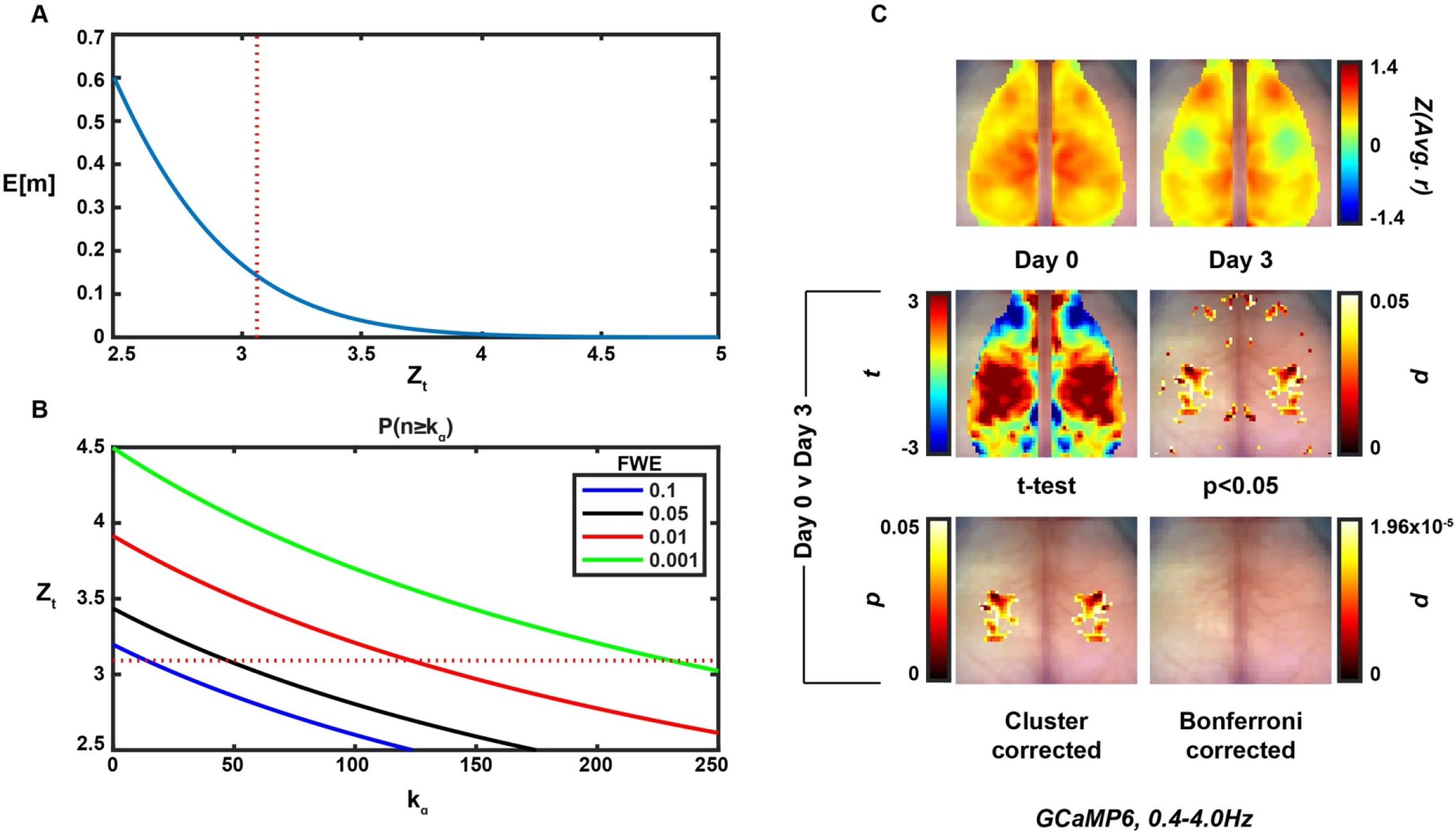
Random field theory determines cluster size thresholding for pixel-wise t-maps. A) The relationship between the pixel-wise false positive rate (determined by Z_t_, dashed red line is Z_t_=3.09 which corresponds to p=0.001) and the expected number of clusters to survive thresholding due to chance (E[m]). B) The relationship between the pixel-wise false positive rate (determined by Z_t_, dashed red line is Z_t_=3.09 which corresponds to p=0.001), the family-wise error rate (here, 0.05) and cluster size needed for significance (*k*_*α*_). C) (top row) Average (N=4) bilateral FC maps pre (left) and three days post (right) photothrombotic stroke to left somatosensory forepaw cortex. (middle row) Pixel-wise paired t-test (left) and thresholded image for pixels with p<0.05 (right). (bottom row) Thresholded image with FWE=0.05 using the cluster size-based method (left) and image with Bonferroni correction for multiple comparisons (right).

**Figure 3:**
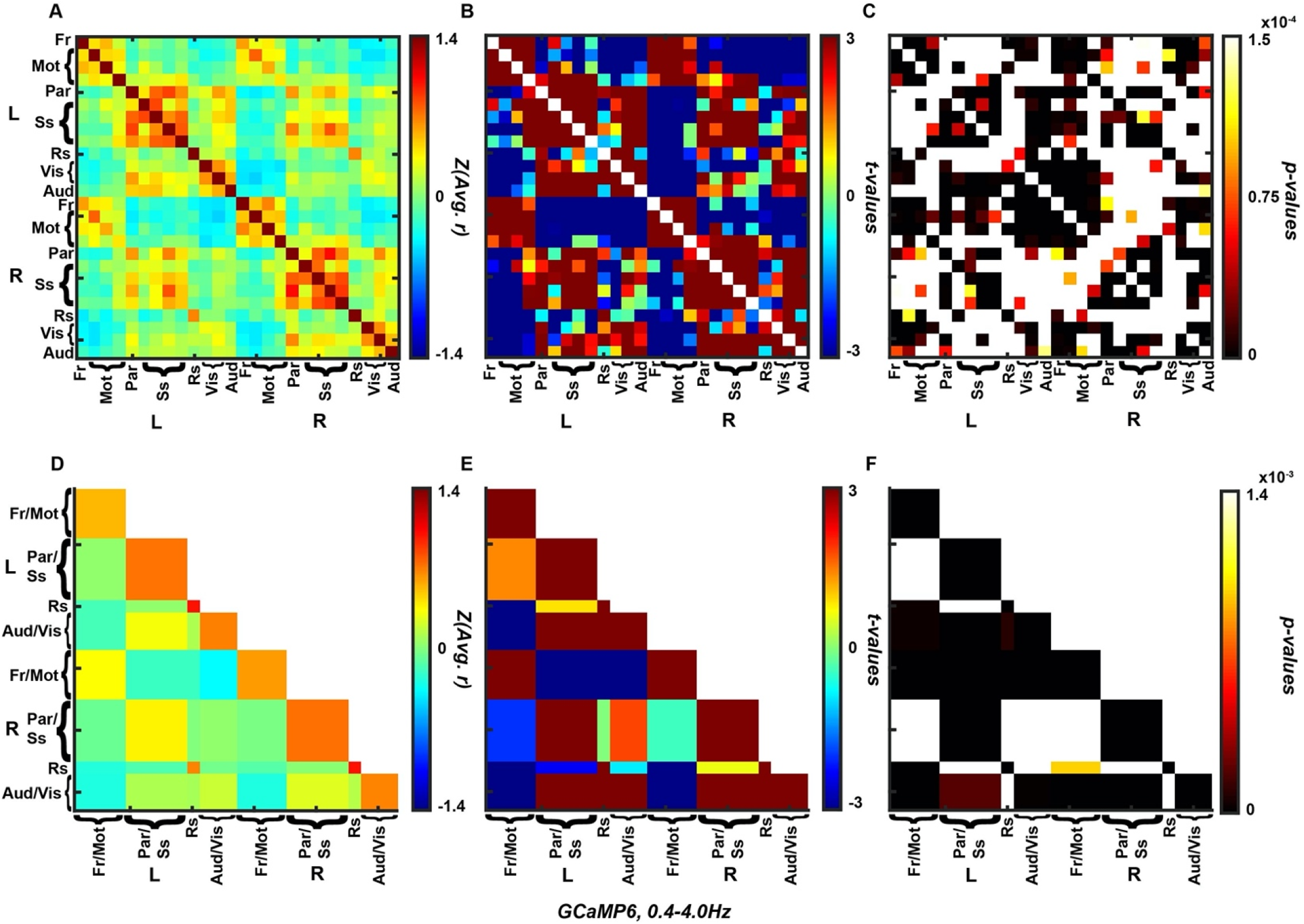
Wild-type mouse FC matrix using traditional and enrichment-based methods. A) Average (N=16) traditional FC matrix. B) Comparison-wise one-sided t-test. C) Matrix thresholded for p-values beneath the Bonferroni threshold for significance (p=1.5×10^−4^). D) Average (N=16) enriched FC matrix. E) Comparison-wise one-sided t-test. F) Matrix thresholded for p-values beneath the Bonferroni threshold for significance (p=1.4×10^−3^).

### Cluster size-based thresholding applied to stroke data

A cluster size-based statistical thresholding method^15,16^ was adapted from the fMRI and DOT literature and used to select clusters of size *k*_*α*_ (expressed in number of pixels) with p<0.05 by the pixel-wise paired t-test method that satisfies the pixel-wise false positive rate (set by Z_t_) and overall family-wise error rate (Figures 2A,B). Using this cluster size cutoff, we were able to compare bilateral FC maps at baseline (N=4, Day 0) and 72 hours post (N=4, Day 3) photothrombotic stroke (Figure 2C). Photothrombosis was induced in left somatosensory cortex which resulted in loss of homotopic FC at Day 3. A pixel-wise t-test was performed and thresholded to only display regions with p<0.05 (note, this map is not corrected for multiple comparisons). Using the cluster-size based threshold (FWE=0.05) we were able to localize a somatosensory anchored deficit. Using the Bonferroni correction for multiple comparisons, no regions survived this stringent cutoff, resulting in no significant differences between Day 0 and Day 3 with this method.

### Matrix enrichment presents easily digestible resting state FC

Mice that had not undergone any experimental manipulations were imaged and FC matrices utilizing all standard seeds in the FOV^23^ were computed (N=16, Figure 3A). A one-sided t-test was performed on each matrix index to isolate significant positive or negative correlations (Figure 3B). A Bonferroni correction for multiple comparisons resulted in a number of significant positive and negative correlations present at rest in non-experimental mice (Figure 3C). To decrease the number of comparisons and improve readability of plots, a data reduction technique was implemented to “enrich” the FC matrices (N=16, Figure 3D). Similarly, a comparison-wise one-sided t-test was performed (Figure 3E) and the matrix was thresholded using a Bonferroni correction for multiple comparisons (Figure 3F).

## Discussion

Wide-field optical imaging (WOI) of mice expressing genetically encoded calcium indicators (GECIs) provides cell-specific, improved temporal resolution recordings of calcium transients across the entire mouse cortex^8–10^. Functional neuroimaging analysis pipelines have been thoroughly developed in the functional magnetic resonance imaging (fMRI) literature^24,25^, however, despite having many similarities, these analytical techniques have not been translated into a comprehensive, user friendly mouse WOI toolbox. Here, we provide a toolbox that pre-processes data according to previous reports^8,20^. We also implement some statistical approaches from the fMRI literature to handle the multiple comparisons problem present in all functional neuroimaging data.

The spatial resolution afforded by techniques such as fMRI or WOI allow neuroscience inquiries with mesoscopic-level spatial specificity^2,9^. However, treating each voxel or pixel as an independent measure will result in a frequently insurmountable threshold when correcting for multiple comparisons (e.g., using the Bonferroni correction). Therefore, there has been significant work done in the fMRI literature to develop more appropriate algorithms to address this^15^. The cluster extent-based thresholding statistical approach operates on the hypothesis that neighboring pixels are likely not independent samples (i.e., a Bonferroni correction for multiple comparisons would be too stringent) and therefore large-grouped differences via independent pixel-wise statistical tests are more likely to represent a significant change somewhere within that cluster. Here, this method is set up to have a family wise error (FWE) rate of 5%, meaning in the collection of thresholded pixel-wise t-tests, there is a 5% chance of having at least one false positive result. An advantage of this technique is it allows for spatial localization of an activation or change in connectivity by considering all the pixels or voxels in the FOV. However, this method only works as well as the initial analysis does in spatially isolating an activation or change, since the rightful conclusion of a cluster surviving thresholding is that a change or activation occurred within that cluster. Here, we translate this method commonly used in three-dimensional fMRI voxel space to two-dimensional WOI pixel space and demonstrate the ability to localize a somatosensory-based deficit three days after a photothrombotic event to left somatosensory cortex (Figure 2C).

Another approach to solving the multiple comparisons problem is through data reduction prior to statistical testing, so the number of statistical comparisons decreases (the light blue boxes in Figure 1C, compared to the cluster size-based method that occurs at the last dark blue box in Figure 1C). One commonly used summary of FC using multiple seeds is by presenting FC matrices (Figure 3A). This way, instead of having multiple FC maps (i.e., one per seed) consisting of as many pixels or voxels in the FOV, each brain region is reduced to one or more seed regions. However, a well sampled FOV can still result in hundreds or thousands of comparisons, making a Bonferroni correction for multiple comparisons a stringent threshold to pass. Here, we propose reducing the data within a common seed x seed FC matrix to have one singular value per cortical region (i.e., per network). Neighboring cortical regions with small surface areas (e.g., Frontal, Parietal, Auditory) were grouped into the neighboring larger functionally similar cortical region to enhance data reduction. Also note that with at least Auditory and Frontal cortex, these cortical regions can often be outside of the FOV^9^. Not only does this matrix methodology still capture inter and intra-cortical dynamics, but the multiple comparisons problem because more feasibly solved by a Bonferroni correction for multiple comparisons (both because the threshold increases by an order of magnitude but also because different brain networks are more likely to be independent samples than neighboring pixels). Shown here, we are able to isolate the strongly positive and negative network correlations across the dorsal cortical surface of the mouse by performing a one-sided t-test at each matrix index (Figure 3). While significant positive and negative cortical correlations are present here with and without data enrichment methods, it is feasible that with a disease model (as in Figure 2C), a less stringent threshold for significance will be necessary to correctly detect an effect.

## Conclusions

This toolbox fills a much-needed gap between the fMRI and WOI data processing communities. Shown throughout are examples of statistical measures that were developed for fMRI being applied to FC calculated with the WOI data. However, the toolbox is also set up to compute multiple types of analyses (detailed in Results), which can all be plotted topographically, statistically compared, and corrected via cluster-based thresholding. The wide distribution and use of this toolbox will greatly aid groups that are hoping to start imaging mouse models of health and disease in order to better understand how brain dynamics might change in humans.

## Acknowledgments

We would like to thank Brian White, Patrick Wright, Adam Bauer, Rachel Rahn, and Hunter Banks who either contributed some iteration of code or test ran software releases to make this project possible. This work was supported by the National Institutes of Health [grant numbers R01NS099429, R01NS090874] awarded to J. P. C., and the National Institute on Aging [grant number F30AG061932] awarded to L. M. B.

